# Atlas-scale single-cell analysis beyond in-memory paradigm with scAtlasPy

**DOI:** 10.64898/2026.09.03.748766

**Authors:** Han Xu, Yangzhan Ye, Senpeng Zhang, Runzhi Xie, Junping Li, Jiadong Lin, Yuxuan Hu, Lin Gao

## Abstract

Single-cell atlases are rapidly outgrowing the memory capacity of standard workstations, challenging the in-memory paradigm underlying mainstream computational ecosystems. Here, scAtlasPy decouples scale of atlas from memory capacity by leveraging the disk-resident computing. It enables full-resolution analysis of a 100-million-cell atlas with only 42.9 GB peak memory, whereas state-of-the-art platforms are limited at 3 million cells with 512 GB memory. scAtlasPy achieves 137,745 cells/s, 10.4× faster than scDataset with 82.6% lower memory usage for random minibatch retrieval. Its extensible architecture offers a flexible platform for diverse atlas-scale analytical tasks, facilitating the discovery of complex cellular heterogeneity and functions in massive cell atlases.

## Main

Single-cell atlases enable high-resolution dissection of cellular heterogeneity across healthy and diseased tissues and organs^1,2^. As atlases expand to millions and even hundreds of millions of cells^3,4^, their scale is increasingly exceeding the memory capacity of computing systems and challenging the in-memory paradigm underlying mainstream single-cell analysis methods^5,6^. This growing mismatch will limit the discovery of novel and complex cellular functions hidden within the increasingly large cell atlases. To address this bottleneck, we developed scAtlasPy, a platform that enables efficient atlas-scale computation beyond in-memory paradigm.

Although efforts have been made to accommodate massive datasets, existing solutions remain constrained by trade-offs in analytical resolution, workflow completeness, and extensibility, and have not fully bridged the fundamental gap between rapidly growing single-cell atlases and the limited memory capacity of conventional computing systems. Within the R ecosystem, Seurat cannot natively handle matrix containing more than 2^3^^1^-1 non-zero entries (**Supplementary Note 1**). It adopts sketch-based analysis^7^ as a practical compromise for atlas-scale analysis, which reduces dataset scale and projects results back to the full atlas. Nevertheless, this approach dilutes analytical resolution, which may depart from the original purpose of large-scale cell atlases: the comprehensive characterization of high-resolution cellular heterogeneity^3^. BPCells^8^ maintains a small memory footprint by retaining data on disk and delivers highly efficient computation for specific operators (i.e., normalization and principal component analysis), yet its operator-oriented strategy is less readily adaptable to newly proposed analytical algorithms and advanced downstream models. Within the Python ecosystem, Scanpy’s backed mode offers on-disk access to atlas-scale data, but provides limited support for the operations required by exploratory single-cell atlas analysis pipelines (**Extended Data Table 1**; **Supplementary Note 2**). scDataset^9^ provides multiple data-indexing strategies based on Scanpy’s backed mode, while lacking a basic data management mechanism for flexible exploratory single-cell atlas analysis workflows. AnnSQL^10^ employs DuckDB as the underlying database engine to facilitate the analysis of atlas-scale datasets. However, its routine analyses rely on direct interaction with the underlying database, requiring expertise in Structured Query Language (SQL) and database systems, and the scalability to large datasets remains limited. These limitations call for a dedicated on-disk platform that moves atlas-scale analysis beyond in-memory limits, supports complete analytical workflows, and provides an extensible foundation for the broader atlas-scale computational ecosystem.

scAtlasPy supports analytical workflows and atlas-oriented computational methods under strict memory constraints. Specifically, scAtlasPy adopts a three-layer architecture comprising a hybrid sparse storage layer, a high-throughput retrieval engine and an open application layer (**Figure 1A, B; Methods**). The storage layer uses a dual-sparse design to balance compact disk representation with efficient on-demand data access (**Extended Data Figure 1; Methods**). The retrieval engine provides standardized interfaces for flexible data loading, querying and random streaming access (**Figure 1C**). Built on this engine, the application layer enables complete atlas-scale analysis workflows under strict memory constraints and enables flexible integration of emerging analytical algorithms and custom models (**Figure 1D**).

**Figure 1.**
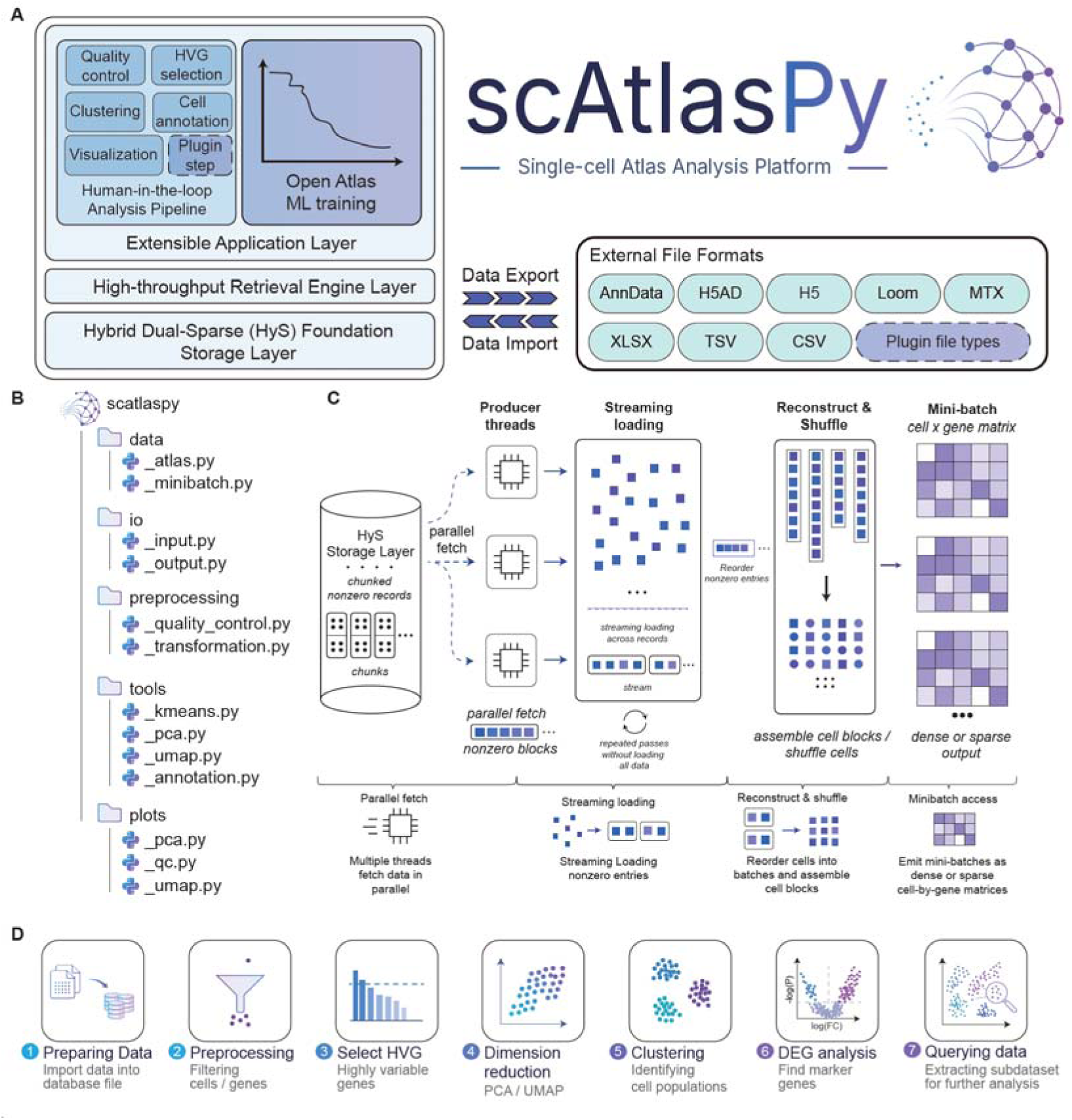
Overview of the scAtlasPy platform. A, Overview of the scAtlasPy architecture. **B,** Modular organization of the scAtlasPy software package. **C,** High-throughput random data streaming for mini-batch generation. **D,** scAtlasPy supports basic single-cell analysis workflow at atlas-scale.

To assess whether scAtlasPy decouples exploratory atlas workflows from physical memory capacity, we benchmarked it against representative memory-efficient single-cell analysis platforms using Tahoe-100M subsamples ranging from 10 thousand to 30 million cells (**Extended Data Table 2**). The benchmark covered a standard exploratory workflow, including quality control, normalization, log transformation, highly variable gene selection, dimensionality reduction and clustering. Across all dataset scales, scAtlasPy completed the entire workflow with peak memory usage never exceeding 27.5 GB, whereas other approaches exceeded available memory, reached time limits or encountered runtime failures as dataset size increased (**Figure 2A; Extended Data Table 3;** Supplementary Table 1-6).

**Figure 2.**
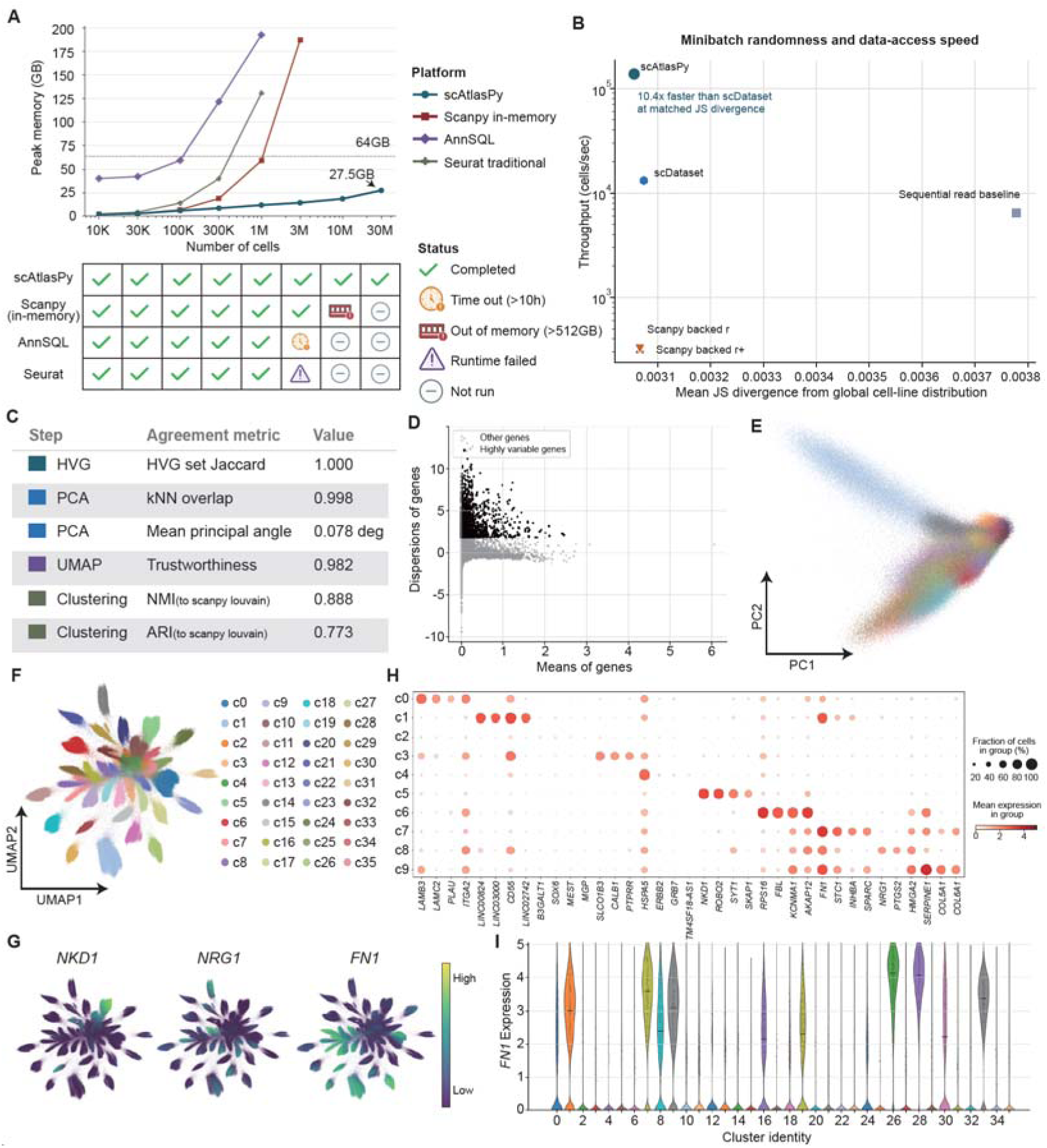
scAtlasPy enables full-resolution atlas-scale exploratory analysis beyond in-memory limits. A, Peak memory and run status for full-resolution workflows on Tahoe100M subsets from 10K to 30M cells. The dashed line marks 64 GB memory. **B,** Mini-batch retrieval speed and randomness on the Tahoe100M plate-1 subset. Randomness is measured by mean Jensen-Shannon divergence from the global cell-line distribution. **C,** Agreement between scAtlasPy and in-memory Scanpy on HLCA core data across HVG, PCA, UMAP and clustering metrics. **D-I**, Representative exploratory outputs from the 100M-cell Tahoe atlas generated using scAtlasPy: **(D)** HVG selection, **(E)** PCA, **(F)** UMAP clustering, **(G)** gene feature plots, **(H)** marker dot plot, and **(I)** marker violin plot. **(E)** and **(F)** are colored by cluster label.

We next assessed whether the streaming architecture of scAtlasPy could provide the data-access performance required by emerging atlas-oriented computational methods. scAtlasPy achieved a random minibatch retrieval throughput of 137,745 cells per second, outperforming the backed mode of Scanpy, a widely adopted single-cell analysis framework, by 425-fold and scDataset, the second-fastest approach in our benchmark, by 10.4-fold (**Figure 2B; Extended Data Figure 2**). Importantly, this scalability did not compromise analytical accuracy. On HLCA^11^, the built-in streaming implementations of highly variable gene selection, PCA, UMAP and Louvain clustering closely reproduced the established Scanpy workflow, achieving an HVG Jaccard score of 1.0, a PCA kNN overlap of 0.998 with a mean principal angle of 0.078, a UMAP trustworthiness score of 0.982 and a clustering NMI of 0.888 (**Figure 2C; Extended Data Figure 3**). Beyond these quantitative comparisons, the resulting embedding representation preserved biologically coherent cellular structure: PCA separated major cell compartments and retained characteristic marker-expression patterns (**Extended Data Figure 4D, E**), while UMAP recovered fine-grained cell-type organization and lineage-associated expression of markers such as *CD3D* (**Extended Data Figure 4G, H**). Notably, coloring the UMAP-embedding by source dataset also revealed pronounced cross-dataset batch effects (**Extended Data Figure 4F**), highlighting substantial technical heterogeneity within the assembled atlas. Downstream differential-expression and marker analyses further resolved cluster-specific transcriptional programs (**Extended Data Figure 4I, J**). These observations illustrate that increasingly large cell atlases pose not only computational challenges but also methodological demands for algorithms capable of resolving complex technical variation directly at atlas scale^3,12^, for which the high-throughput streaming interfaces of scAtlasPy provide an extensible computational foundation.

To demonstrate that scAtlasPy scales to extremely large atlases under constrained computational resources, we applied its built-in workflow to full-resolution exploratory analysis of the Tahoe-100M atlas on a laptop with 64 GB of memory. The analysis proceeded from highly variable gene selection (**Figure 2D**) and PCA (**Figure 2E**) to clustering analysis and UMAP visualization (**Figure 2E,F**), resolving 36 transcriptionally distinct clusters across the full atlas. Gene feature maps further revealed heterogeneous expression patterns of *NKD1*, *NRG1* and *FN1* across these clusters (**Figure 2G**), while cluster-resolved dot plots and violin plots characterized distinct marker-expression programs (**Figure 2H, I**). Besides quality control, differential-expression and marker analyses further demonstrated that the workflow could extend beyond dimensionality reduction and clustering to downstream gene- and cluster-level interpretation at full atlas scale (**Extended Data Figure 5**). The complete workflow finished in 7.68 hours with a peak memory usage of 42.86 GB, remaining within the 64 GB memory budget throughout all analytical stages (**Extended Data Figure 6A, B**). These results demonstrate that full-resolution exploratory and downstream analysis of 100-million-cell atlases can be achieved under strictly constrained memory resources in the framework of scAtlasPy.

Notably, the primary goal of scAtlasPy is not to replace existing single-cell analysis platforms with another universal toolkit, but to provide an on-disk foundation that enables seamless integration of atlas-scale computational analysis, fostering the continued evolution of the atlas-scale analysis ecosystem^13^ (**Extended Data Figure 7**). With the rapid expansion of single-cell atlas datasets, the growing mismatch between massive data volume and limited computational resources has become a major bottleneck in single-cell analysis. Existing mainstream tools, such as Seurat and Scanpy, are well suited for routine analyses of small- to medium-sized datasets, whereas other specialized frameworks can substantially accelerate computation when enough GPU memory is available^14^. In contrast, scAtlasPy is designed to support full-resolution analysis of ultra-large cell atlases under strictly constrained memory conditions.

A key advantage of scAtlasPy lies in its highly available random data streaming capability, which can facilitate the development of novel atlas-oriented analytical algorithms by providing stable and efficient data stream support. To enhance usability and accessibility, scAtlasPy also supports the import and export of multiple popular data formats (e.g., h5ad) and offers application programming interfaces (APIs) with a Scanpy-like design, minimizing the learning curve for researchers familiar with established single-cell analysis workflows. Furthermore, the underlying storage architecture of scAtlasPy is designed to be language- and platform-agnostic, avoiding dependence on a specific programming environment and enabling future interoperability across major single-cell analysis ecosystems. This design makes it possible to extend the same disk-resident data infrastructure to R-based workflows without redesigning the underlying atlas representation. An important next step is the development of an optimized R-compatible retrieval engine, which would allow R-based analytical methods to access the same atlas-scale data infrastructure and facilitate interoperability across programming environments.

Overall, scAtlasPy establishes a disk-resident and streaming-enabled foundation for atlas-scale single-cell analysis. The significance of this architecture extends beyond its current implementation, enabling the broader atlas-scale computational ecosystem to evolve beyond the in-memory paradigm and supporting future methods that can uncover increasingly complex cellular heterogeneity and functions from massive cell atlases.

## Methods

### Three-layer Architecture of scAtlasPy

scAtlasPy is implemented as a hierarchical framework composed of three coordinated layers: a hybrid foundational storage layer, a high-throughput retrieval engine layer, and an extensible application layer. This design separates physical data persistence, scalable data access, and analytical computation, allowing large-scale single-cell atlas workflows to be executed without materializing the full expression matrix in memory. Its modular architecture further provides a unified foundation for integrating new atlas-scale analytical methods within the disk-resident streaming framework.

### Hybrid Storage Layer

At the foundation of scAtlasPy is a hybrid persistent storage layer that leverages DuckDB as an embedded database backend while organizing atlas-scale single-cell omics data through a customized relational schema. Each atlas is represented by an Atlas object, which manages a single on-disk *.sasql* database file and provides the database connection, query execution, and lifecycle management. Single-cell datasets are converted from AnnData-compatible inputs into a relational representation consisting of cell metadata, gene metadata, sparse expression tables, and embedding or model-result tables.

The expression matrix adopts a dual-sparse storage design. The raw count matrix is encoded by a combination of compressed sparse row (CSR) and Coordinate list (COO) format, where non-zero expression values are stored in *X_HyS*, and row pointer information is stored in *X_HyS_indptr*. Each non-zero entry records the corresponding cell ID, gene ID, and expression value, and the pointer array enables efficient reconstruction of cell-level sparse vectors. This design integrates the advantages of both CSR and COO sparse formats (**Extended Data Figure 1**), delivering more flexible data retrieval strategies than any single sparse storage scheme while maintaining superior compression efficiency.

Besides expression values, scAtlasPy stores structured atlas annotations in relational tables. Cell-level metadata are stored in *obs*, gene-level metadata in *var*, cell embeddings such as PCA or UMAP in *obsm_\** tables, gene loadings in *varm_\** tables, and algorithmic statistics or parameters in *uns_\** tables. This schema mirrors the logical organization of *AnnData* while replacing in-memory containers with a queryable, persistent database representation.

To ensure full traceability of exploratory analysis operations, scAtlasPy first generates filtering flags in the *obs* and *var* tables for every data-filtering procedure. Upon completion of quality control, feature selection, and data transformation, the framework maintains a filtered and reindexed sparse matrix representation. Filtered cells and genes are remapped to contiguous *filter_cell_id* and *filter_gene_id*, and a derived sparse table *X*_*HyS_filtered* is constructed to support downstream minibatch analysis. This strategy eliminates empty rows and columns after filtering, and enables the reconstruction of both dense and sparse minibatches from the refined analytical matrix.

### High-Throughput Retrieval Engine Layer

The retrieval engine provides standardized on-disk data access interfaces for database-resident atlases. After import, scAtlasPy stores the sparse expression matrix in DuckDB using a CSR-inspired representation, with nonzero expression values stored separately from per-cell pointer arrays. This layout enables efficient reconstruction of selected cells or minibatches without scanning or materializing the full cell-by-gene matrix.

During minibatch retrieval, the engine reads only the required nonzero records from DuckDB, reconstructs the corresponding sparse cell-by-gene block, and converts it to a dense array only when requested by downstream algorithms. The retrieval interface supports both single-pass and multi-pass access. In single-pass mode, cells are streamed through the atlas once. In multi-pass mode, the engine repeatedly scans the atlas while maintaining a shuffle buffer that generates randomized minibatches across epochs without loading the complete dataset into memory.

To maximize throughput, scAtlasPy adopts a multithreaded producer-consumer architecture. Producer threads asynchronously issue database queries for upcoming minibatches, while the consumer reconstructs batches in order and returns standardized dense arrays to downstream analytical modules. This design overlaps database I/O with computation, reducing retrieval latency while providing a unified data interface for streaming PCA, minibatch clustering, UMAP training, and machine-learning workflows.

Together, these components establish the retrieval engine as the bridge between the physical DuckDB storage layer and downstream numerical analysis. It provides both minibatch-level and cell-selection-level access while keeping memory usage bounded by the requested batch size, selected features, and shuffle-buffer configuration rather than by the total number of cells in the atlas.

### Extensible Open Application Layer

The top layer implements user-facing analytical modules in a Scanpy-like namespace organization: *scatlaspy.pp* for preprocessing, *scatlaspy.tl* for computational tools, *scatlaspy.pl* for visualization, and *scatlaspy.io* for input and output. These modules operate through the standardized retrieval layer rather than directly depending on in-memory objects. As a result, the single-cell workflows of scAtlasPy can be executed over database-resident atlases.

The preprocessing module provides database-backed cell and gene filtering, quality-control metric calculation, library-size normalization, logarithmic transformation, scaling, and highly variable gene selection. The tool module implements atlas-scale analytical methods, including streaming PCA, distilled Louvain clustering, minibatch K-means clustering, teacher-student UMAP, and marker-gene ranking. Results are written directly back to the same atlas database as metadata columns, embeddings, loadings, cluster labels, or unstructured result tables, enabling downstream analysis and visualization without regenerating intermediate objects.

The plotting module reads directly from database-resident metadata, embeddings, and expression tables to generate common atlas visualizations, including QC plots, highly expressed gene summaries, HVG plots, PCA and UMAP embeddings, feature plots, violin plots, dot plots, volcano plots, cluster-size distributions, and marker-gene summaries. The IO module supports import from AnnData-compatible formats and export to .h5ad, preserving interoperability with the broader single-cell ecosystem.

Because the application layer communicates with the database through standardized retrieval and table-writing interfaces, new algorithms can be integrated as plugins or custom analytical modules. Such extensions can consume minibatches, sparse tables, metadata tables, or existing embeddings, and persist their outputs using the same *obs, var, obsm, varm,* and *uns* conventions. This design enables scAtlasPy to support both standard single-cell analytical workflows and emerging atlas-oriented computational methods within a unified disk-resident framework.

### Randomness of mini-batch retrieval

scAtlasPy achieves randomized minibatch retrieval through two complementary stages: atlas import and downstream data access. During import, cells are not written to the atlas database in their original order. Instead, scAtlasPy reads contiguous blocks of cells from the input dataset, aggregates multiple blocks into a memory-bounded import window, randomly permutes the cells within the window, and writes the shuffled cells together with the corresponding sparse expression records and CSR-inspired pointer arrays into the database. Consequently, the database-resident atlas is partially randomized before downstream analysis begins, reducing potential ordering biases inherited from the input dataset.

To further improve mixing across neighboring regions of the dataset, scAtlasPy employs a rolling shuffle strategy during import. At each import window, only a portion of the shuffled cells is written immediately, while the remainder is carried forward and mixed with cells from the next window. This carry-over mechanism propagates randomness across consecutive import windows rather than restricting it to individual windows. The import window size is determined by the configured database memory limit, allowing the degree of randomization to scale with available memory without changing downstream interfaces.

During minibatch retrieval, scAtlasPy reconstructs cell-by-gene matrices directly from the database-resident sparse representation. A read index stores filtered cells and genes in CSR-inspired order, enabling each minibatch to be reconstructed from contiguous sparse records. For iterative algorithms, scAtlasPy provides a multi-pass retrieval mode in which the atlas is scanned repeatedly while an internal shuffle buffer continuously mixes minibatches across the data stream before they are returned to the algorithm. This design prevents minibatch composition from becoming fixed across successive passes, providing sustained stochasticity for iterative optimization.

For applications in which each cell should be visited only once, scAtlasPy also supports single-pass retrieval. In this mode, minibatches are generated from a single traversal of the read index, preserving the same memory-bounded sparse reconstruction pathway while avoiding repeated scans of the atlas. Single-pass retrieval is suitable for throughput benchmarking and deterministic analyses, whereas multi-pass retrieval supports iterative algorithms that benefit from repeated randomized exposure to the data.

Together, import-time rolling randomization and retrieval-time streaming shuffling provide effective minibatch mixing while preserving the performance advantages of on-disk analysis. Both the input dataset and the database are accessed predominantly through large sequential or near-sequential reads, while only the current minibatch and shuffle buffer reside in memory. This design eliminates the need for global in-memory permutation of cell indices, enabling randomized minibatch access for atlas-scale datasets under strict memory constraints.

### Data import and export

scAtlasPy supports flexible data exchange with common single-cell data formats, including *AnnData .h5ad*, *10x Genomics .h5*, *Matrix Market .mtx*, *Loom*, and in-memory *AnnData* objects. This broad compatibility allows users to initialize an atlas from standard single-cell datasets while retaining interoperability with existing analysis ecosystems.

During data import, input datasets are incrementally converted into persistent DuckDB-backed atlas databases managed by an *Atlas* object. The importer reads AnnData-compatible *.h5ad* files in backed mode and, in the default randomized import mode, processes contiguous blocks of cells, converts expression values to count scale when necessary, mixes cells using a rolling shuffle-window strategy, and incrementally writes them to the database. Cell metadata are stored in the *obs* table, gene metadata in the *var* table, and sparse expression values in a CSR-inspired layout. Nonzero expression records are stored in *X_HyS_data*, while cumulative per-cell row pointers are stored in *X_HyS_indptr*, enabling efficient reconstruction of cell-wise sparse vectors during downstream analysis without materializing the full matrix.

For export, scAtlasPy writes Atlas databases back to AnnData-compatible *.h5ad* files. The exporter reconstructs the sparse expression matrix by resolving the requested expression representation, rebuilding the CSR pointer array, and writing the sparse matrix in batches. Cell and gene metadata are exported as AnnData-compatible dataframes, while stored *obsm_\** and *varm_\** tables are restored to their corresponding *AnnData* slots. This design preserves interoperability with Scanpy and other AnnData-compatible tools without constructing a full in-memory *AnnData* object during export.

### Streaming PCA

scAtlasPy implements PCA using a streaming randomized algorithm^15^ over the on-disk atlas. After feature scaling, the expression matrix is accessed through the minibatch retrieval engine, allowing principal components to be estimated incrementally from successive minibatches without materializing the full cell-by-gene matrix in memory. The learned loading vectors are then used in a second batched pass to project all cells into the low-dimensional space, and the resulting PCA coordinates are written back to the atlas as an *obsm* table. This implementation keeps memory usage primarily dependent on the number of genes, requested components, and minibatch size rather than on the total number of cells.

### UMAP

Classical UMAP becomes increasingly expensive at atlas scale because nearest-neighbor graph construction and iterative layout optimization must be performed over all cells. To enable scalable visualization, scAtlasPy implements a teacher–student UMAP strategy. A conventional UMAP model is first trained on representative cells from the atlas to provide reference low-dimensional embeddings. A lightweight neural network is then trained to learn this embedding and subsequently applied to the entire atlas through minibatch inference. The predicted coordinates are written back to the atlas as the default UMAP embedding. By distilling the low-dimensional structure learned by conventional UMAP into a scalable predictor, this strategy enables atlas-scale visualization while preserving the characteristic organization of the original embedding.

### Clustering

scAtlasPy provides two scalable clustering strategies. The first is minibatch K-means, which consumes dense minibatches from an existing low-dimensional representation, such as PCA or UMAP, and updates cluster centroids incrementally. This approach is fast, memory efficient, and naturally compatible with atlas-scale datasets because it never requires constructing a full cell-cell graph. However, K-means partitions the embedding space using centroid-based decision boundaries and may not fully capture graph-based community structure.

To better approximate conventional graph-based clustering, scAtlasPy also implements a distilled Louvain strategy. A conventional Louvain model first serves as the teacher, assigning graph-based community labels from a representative subset of cells in the low-dimensional embedding. A lightweight classifier is then trained to distill the teacher’s clustering rule directly from the embedding representation. Once trained, the student predicts cluster labels for the entire atlas through minibatch inference, and the resulting labels are written back to the atlas metadata, by default as *scatlas_cluster*. By distilling graph-based community assignments into an efficient classifier, this strategy avoids constructing a full nearest-neighbor graph while preserving Louvain-like clustering behavior at atlas scale.

### scAtlasPy implementation

scAtlasPy is implemented as a Python package with a modular namespace structure designed to separate data management, preprocessing, computational analysis, visualization, and input/output operations. The package exposes a familiar user interface through *scatlaspy.pp, scatlaspy.tl, scatlaspy.pl, and scatlaspy.io,* while the underlying database-resident atlas is managed by the central *Atlas* class.

The Atlas class provides the primary abstraction for a scAtlasPy dataset. Each Atlas instance is associated with a named on-disk *.sasql* DuckDB database and manages database creation, connection handling, SQL execution, and query operations. Analytical functions receive an Atlas object as input and operate directly on its database connection, allowing intermediate results to remain persistent throughout the workflow.

The package is organized into five main modules. The data module implements the Atlas object, filtered sparse-matrix reconstruction, and multithreaded minibatch retrieval. The io module provides dataset import and export routines, including conversion from AnnData-compatible inputs into DuckDB tables and streaming export back to *.h5ad*. The preprocessing module implements database-backed quality control, filtering, normalization, logarithmic transformation, scaling, and highly variable gene selection. The tools module contains downstream analytical algorithms, including streaming randomized PCA, teacher-student UMAP, distilled graph clustering, and marker-based annotation. The plots module provides visualization functions that query database-resident metadata, embeddings, and expression values.

Most computational procedures are implemented as a combination of SQL operations and batched numerical routines. Operations that can be expressed as aggregations, joins, updates, or table transformations are executed directly in DuckDB, with memory limits and step-specific execution settings used to control temporary memory usage. Algorithms requiring dense numerical input consume data through the minibatch retrieval engine, which reconstructs only the required batches from the sparse database representation.

For dimensionality reduction, scAtlasPy uses a streaming randomized PCA backend. Rather than materializing the full cell-by-gene matrix or full projected sketch in memory, the method scans centered minibatches from the on-disk atlas and accumulates compact cross-product matrices needed to estimate the PCA subspace. The final PCA embeddings are then written back to the atlas in batches. For graph-style clustering at atlas scale, scAtlasPy uses a distilled clustering strategy: a graph clustering teacher is fitted on a representative subset in low-dimensional space, and a lightweight student model assigns labels to the full dataset by batched prediction. This design avoids constructing a full cell-cell graph for all cells while preserving graph-clustering-like behavior for large atlases.

scAtlasPy stores analytical outputs back into the same atlas database using AnnData-inspired conventions. Cell-level results such as QC labels, filtering flags, cluster assignments, and total-count statistics are written to obs; gene-level statistics and feature-selection results are written to var; cell embeddings are stored in *obsm_\**; gene loadings are stored in *varm_\**; and model parameters or diagnostic summaries are stored in *uns_\** tables. This persistent result model allows subsequent modules to reuse previous computations without repeated data loading or recomputation.

The implementation is intentionally extensible. New analytical functions can be added by accepting an Atlas object, accessing standardized database tables or minibatch iterators, and writing results back using the same table conventions. This design allows scAtlasPy to support both built-in single-cell workflows and external atlas-scale machine learning methods while maintaining a consistent storage and execution model.

### Benchmarking

We benchmarked representative single-cell analysis workflows to evaluate scalability, peak memory usage, and minibatch data-loading performance on large-scale single-cell RNA-seq data. All systematic benchmarks were performed using the Tahoe 100M dataset. To generate reproducible benchmark inputs, we selected plates 1-5 and created nested cell subsamples containing 10,000, 30,000, 100,000, 300,000, 1,000,000, 3,000,000, 10,000,000, and 30,000,000 cells. Cells were sampled proportionally from each plate using a fixed random seed of 1. All subsampled datasets retained the original 62,710-feature gene expression space shared across plates 1-5 and were stored as gzip-compressed *h5ad* files. The sparse expression matrices contained 14.2 million to 42.8 billion nonzero entries across the 10k to 30M cell subsets.

For each platform, we evaluated memory usage and runtime using a series of Tahoe100M subsampled datasets. Each platform was tested with a matched exploratory workflow where supported, including data access or import, gene and cell filtering, quality-control metric calculation, library-size normalization, logarithmic transformation, highly variable gene selection, scaling, PCA, and clustering. For scAtlasPy, all steps were executed on the database-resident atlas using its on-disk workflow. For Scanpy and Seurat, the corresponding execution modes supported by each platform were used. Platform failures, including out-of-memory errors, timeouts, unsupported operations, and runtime errors, were recorded, and benchmarking for a platform was terminated once it first failed at a given dataset size. Because Scanpy backed mode does not support a complete exploratory workflow, it was excluded from the workflow-level memory and runtime benchmarking. To characterize its current capabilities, we instead evaluated operation-level support under both backed="r" and backed="r+" using the 10k-cell subset, recording the status of each operation as supported, unsupported, runtime error, or out-of-memory.

We evaluated mini-batch loading performance as a separate benchmark using the Tahoe100M plate-1 subset containing 5,481,420 cells after preparation. This benchmark was performed on the Tahoe100M plate-1 subset containing 5,481,420 cells after preparation, using a mini-batch size of 2,048 cells. After 20 warmup mini-batches, throughput was measured over 2,000 mini-batches, corresponding to 4.10 million retrieved cells per method. We reported mini-batches per second, cells per second, total runtime including warmup, and peak RSS measured across the main Python process and descendant worker processes. Mini-batch randomness was evaluated by comparing the cell-line label distribution within each mini-batch against the global cell-line distribution using Jensen-Shannon divergence, where lower values indicate better agreement with the full-dataset composition. We chose three baseline methods that enable on-disk data manage, including standard backed mode Scanpy, scDataset, and AnnSQL. For the scDataset, we used block-shuffled sampling with BlockShuffling, block_size=4, fetch_factor=16, num_workers=12, and seed=1. For AnnSQL, we wrote a convenience SQL code to retrieval mini-batch. The disk storage cost of AnnSQL and scAtlasPy was also compared.

We evaluated analytical accuracy using the HLCA core dataset, which can be processed by both scAtlasPy and a conventional in-memory Scanpy workflow. Scanpy was used as the reference implementation. Both workflows performed library-size normalization, log transformation, highly variable gene selection, scaling, PCA, neighborhood construction, UMAP, and graph-based clustering under matched analysis settings where applicable. For HVG selection, we compared the selected gene sets between scAtlasPy and Scanpy using Jaccard similarity. For PCA, we compared the resulting low-dimensional representations using k-nearest-neighbor overlap and principal angles between the PCA loading subspaces, thereby assessing both neighborhood preservation and subspace consistency. For UMAP, we evaluated whether the embedding preserved the PCA-space neighborhood structure using trustworthiness. For clustering, scAtlasPy distilled Louvain clustering was compared with Scanpy Louvain clustering using normalized mutual information and adjusted Rand index. Together, these metrics assessed whether scAtlasPy reproduced the key analytical outputs of a conventional in-memory workflow while operating through its on-disk execution engine.

### Memory and Time Measurement

Peak memory usage was quantified using resident set size (RSS). In the scaling benchmark, Python-based platforms were monitored with a background sampler that reads RSS from Linux /proc, and R-based Seurat runs were monitored by a lightweight forked sampler tracking the parent R process. The default sampling interval was 0.2 s. Runtime was measured as wall-clock elapsed time for each workflow step. Each dataset-level run was limited to 10 hours and 512 GB RSS. Runs exceeding the time limit were recorded as timeouts, and runs exceeding the memory limit were recorded as memory failures; in either case, larger dataset sizes for the same platform were not continued. In the minibatch retrieval benchmark, where some methods use worker processes for data loading, peak memory was measured as the combined RSS of the main Python process and descendant worker processes using “memory_profiler” with child-process inclusion enabled.

### Experimental Environments

All benchmarks were run on an Ubuntu 22.04.5 LTS server with Linux kernel 6.8.0-111-generic, two Intel Xeon Gold 6530 CPUs, 128 logical CPU cores, approximately 2 TiB RAM, and a 7.68 TB Solidigm SB5PH27X076T NVMe solid-state drive for dataset I/O. The Seurat benchmark used R 4.5.3, Seurat 5.5.0, SeuratObject 5.4.0, Matrix 1.7.5, reticulate 1.46.0, future 1.70.0, and future.apply 1.20.2. The Scanpy and scDataset benchmarks used Python 3.10.12, Scanpy 1.11.5, AnnData 0.11.4, NumPy 2.2.6, SciPy 1.15.3, pandas 2.3.3, scikit-learn 1.7.1, numba 0.65.1, psutil 7.2.2, PyTorch 2.12.1, and scDataset 0.3.0.

The 100M-cell laptop experiment was run on a MacBook Pro running macOS 26.5.2, equipped with an Apple M5 Pro chip with 18 CPU cores, 64 GB unified memory, and an internal 4 TB Apple SSD AP4096Z NVMe solid-state drive for dataset I/O. The scAtlasPy workflow used Python 3.10.20, scAtlasPy 0.1.0, DuckDB 1.5.4, AnnData 0.11.4, NumPy 2.2.6, SciPy 1.15.3, pandas 2.3.3, scikit-learn 1.7.2, numba 0.66.0, Scanpy 1.11.5, and PyTorch 2.13.0.

## Data availability

All data used in this study were obtained from publicly available database. The HLCA was downloaded from https://cellxgene.cziscience.com/collections/6f6d381a-7701-4781-935c-db10d30de29 <u>3</u> on March 24, 2025. The Tahoe 100M atlas was downloaded from https://console.cloud.google.com/storage/browser/arc-institute-virtual-cell-atlas/tahoe100M/2025-02-25/h5ad on April, 25, 2026.

## Code availability

The implementation of scAtlasPy, all analysis scripts, and experimental results are publicly available. The source code is hosted on GitHub at https://github.com/scAtlasPy/scAtlasPy, and scAtlasPy can be installed directly from PyPI using pip install scatlaspy. Experimental data and results are available through Figshare(https://doi.org/10.6084/m9.figshare.33144860).

## Author contributions

H.X. conducted background research, conceptualized the study, and designed the overall architecture of the platform. H.X. and Y.Y. implemented the Python package and optimized performance bottlenecks. S.Z. collected the data and wrote the software documentation and user manual. H.X., Y.Y., S.Z., R.X. and J.P.L. discussed and refined the implementation strategy. H.X. and Y.Y. performed data visualization. The study was supervised by L.G., Y.H. and J.D.L., with L.G. and Y.H. providing funding support. J.D.L. contributed to the interpretation of the study and refinement of the manuscript. The original manuscript was drafted by H.X. All authors reviewed and edited the manuscript and approved the final version for publication.

## Acknowledgements

We thank Shi-Hua Zhang from the Academy of Mathematics and Systems Science, Chinese Academy of Sciences, for valuable suggestions; Mengmeng Yin, Yiheng Wang and Jianrui Liu from Xidian University for constructive comments and suggestions; and the Xi’an Key Laboratory of Computational Bioinformatics for its support.

This work was supported by the National Natural Science Foundation of China (NSFC) grants No. 62550005, No. 62132015, and No. U22A2037 to Lin Gao; a NSFC grant No. 62422211 and a Scientific Research Innovation Capability Support Project for Young Faculty No. SRICSPYF-ZY2025003 to Yuxuan Hu.

## Extended Data Figures

**Extended Data Figure 1.**
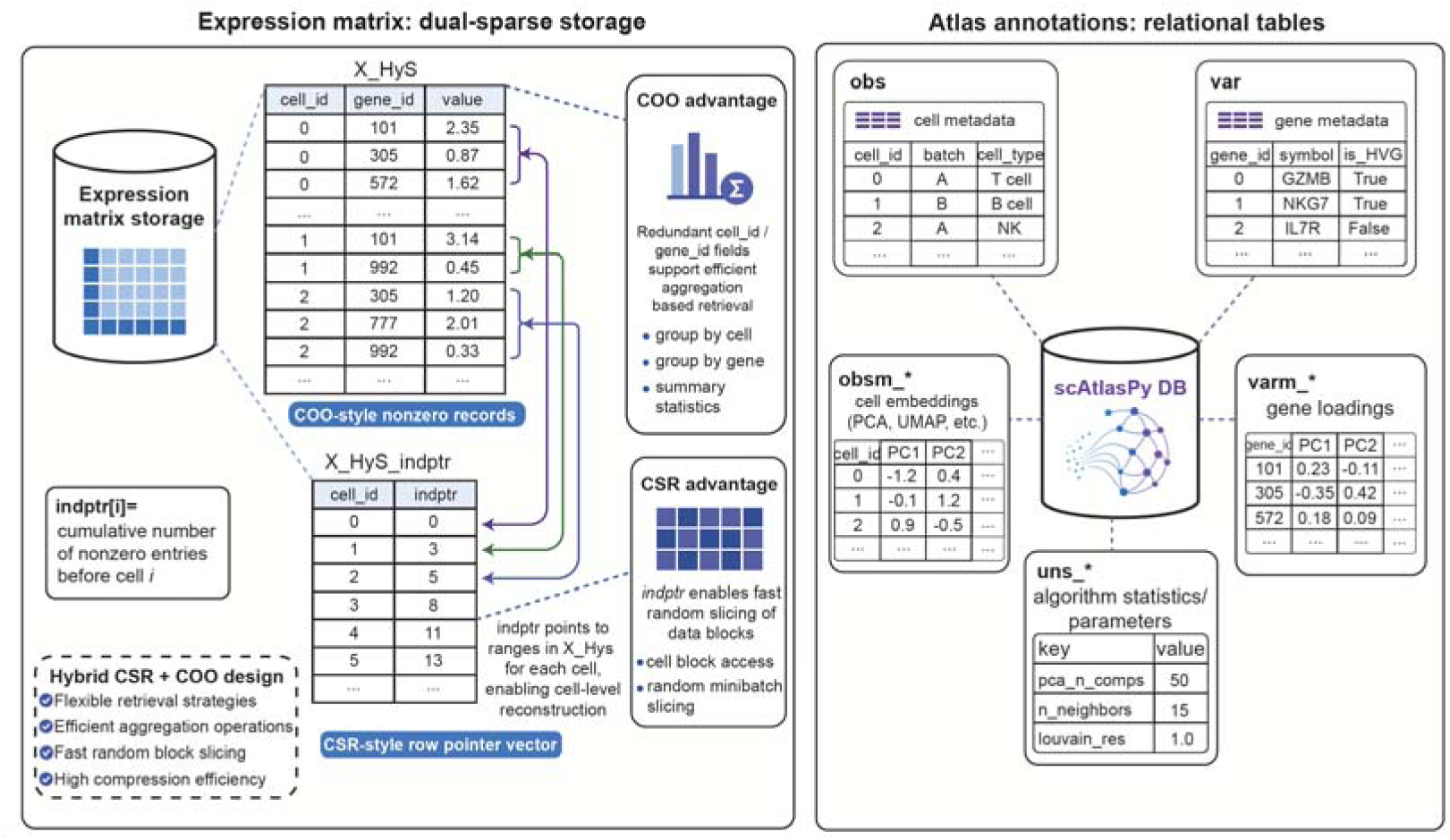
Database architecture of scAtlasPy. Hybrid CSR–COO storage enables efficient expression matrix access (left panel). Relational tables organize atlas annotations and analysis results (right panel).

**Extended Data Figure 2.**
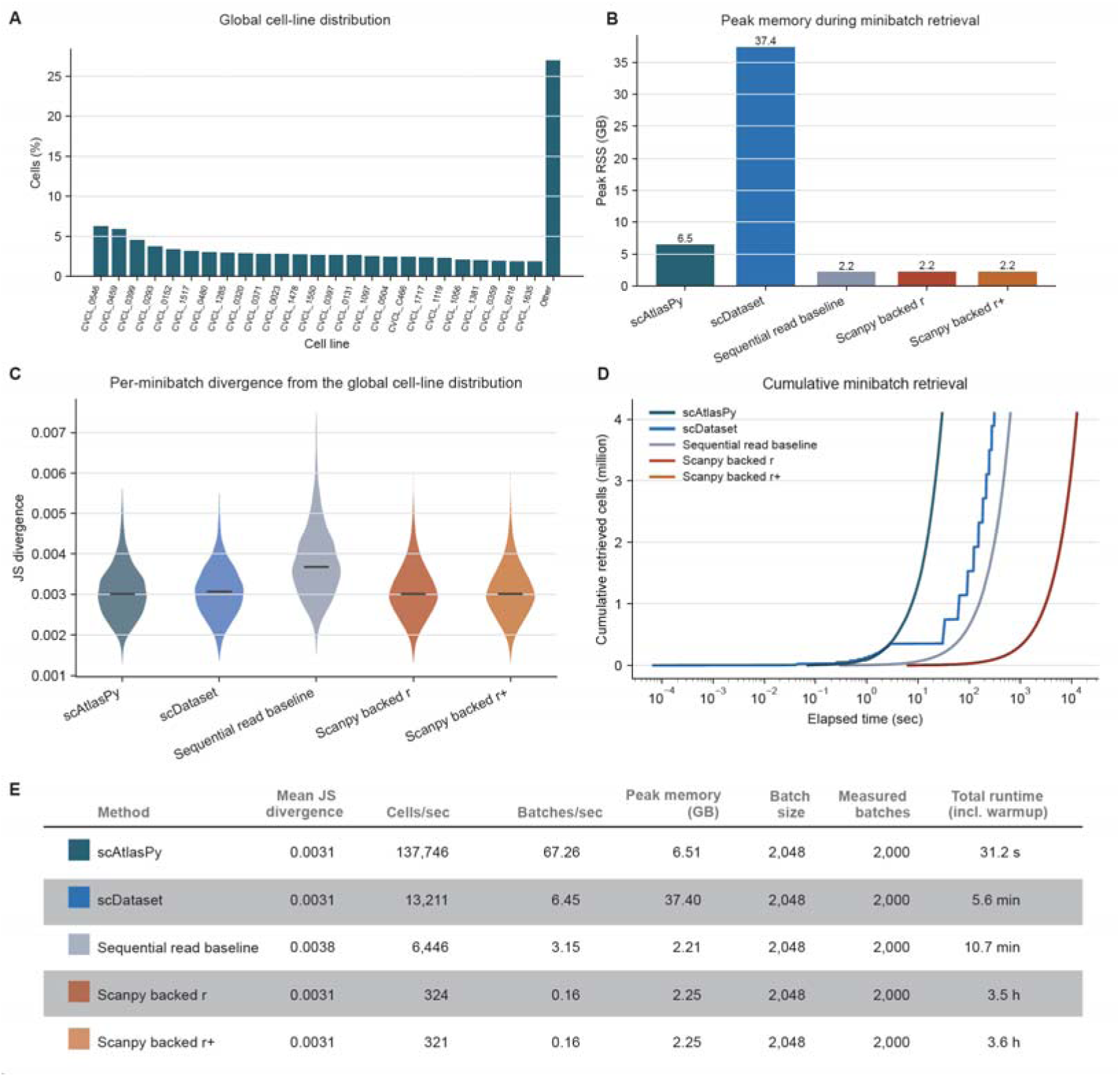
Mini-batch retrieval speed, memory and mixing. A, Global cell-line composition of the Tahoe100M plate-1 subset used for the mini-batch benchmark. **B,** Peak RSS during mini-batch retrieval, including the main Python process and descendant worker processes. **C,** Per-mini-batch Jensen-Shannon divergence from the global cell-line distribution. Lower values indicate better mini-batch mixing. **D,** Cumulative retrieved cells over elapsed time for each method. **E,** Summary of mini-batch retrieval performance. Runtime includes 20 warmup mini-batches followed by 2,000 measured mini-batches with a batch size of 2,048 cells.

**Extended Data Figure 3.**
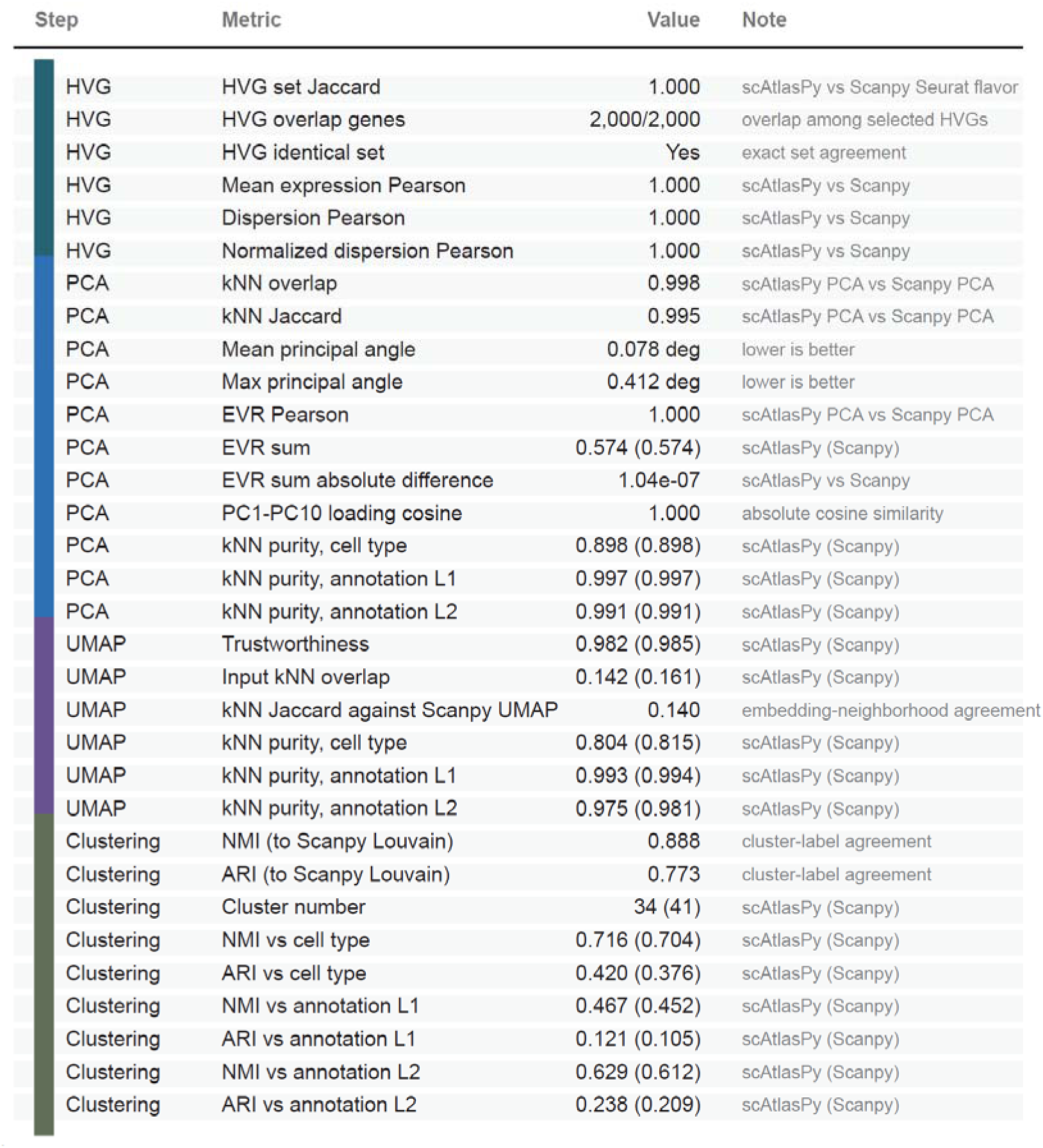
Full summary of agreement between scAtlasPy and the in-memory Scanpy reference workflow on the HLCA core dataset.

**Extended Data Figure 4.**
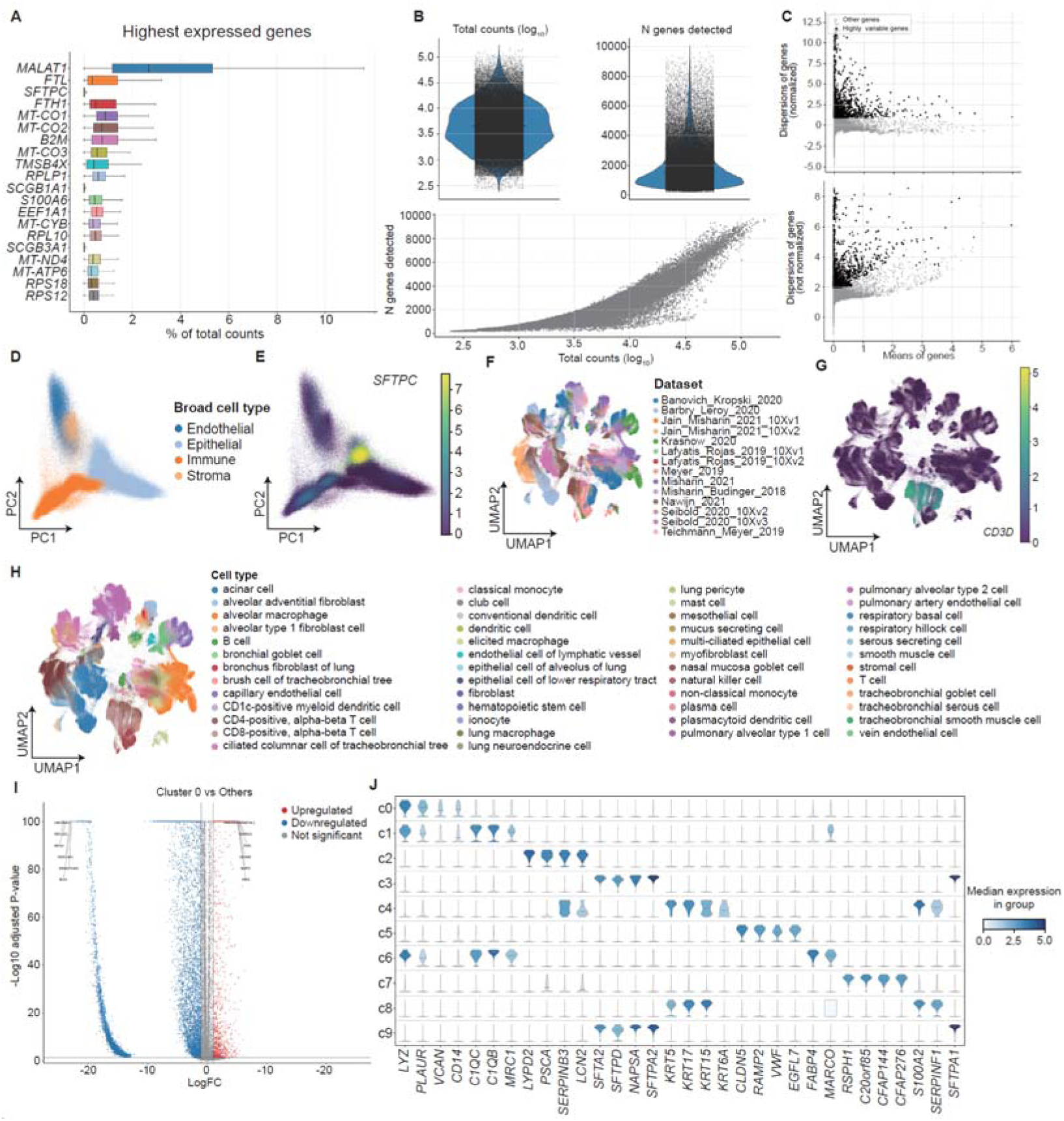
HLCA core exploratory analysis with scAtlasPy. Example scAtlasPy workflow on the HLCA core dataset, including (**A**) highest expressed genes, (**B**) QC summaries, (**C**) HVG selection before and after normalization, (**D,E**) PCA visualizations by broad cell type and marker expression, (**F-H**) UMAP visualizations by dataset, marker expression and cell type, (**I**) differential-expression volcano plot and (**J**) cluster marker violin plot.

**Extended Data Figure 5.**
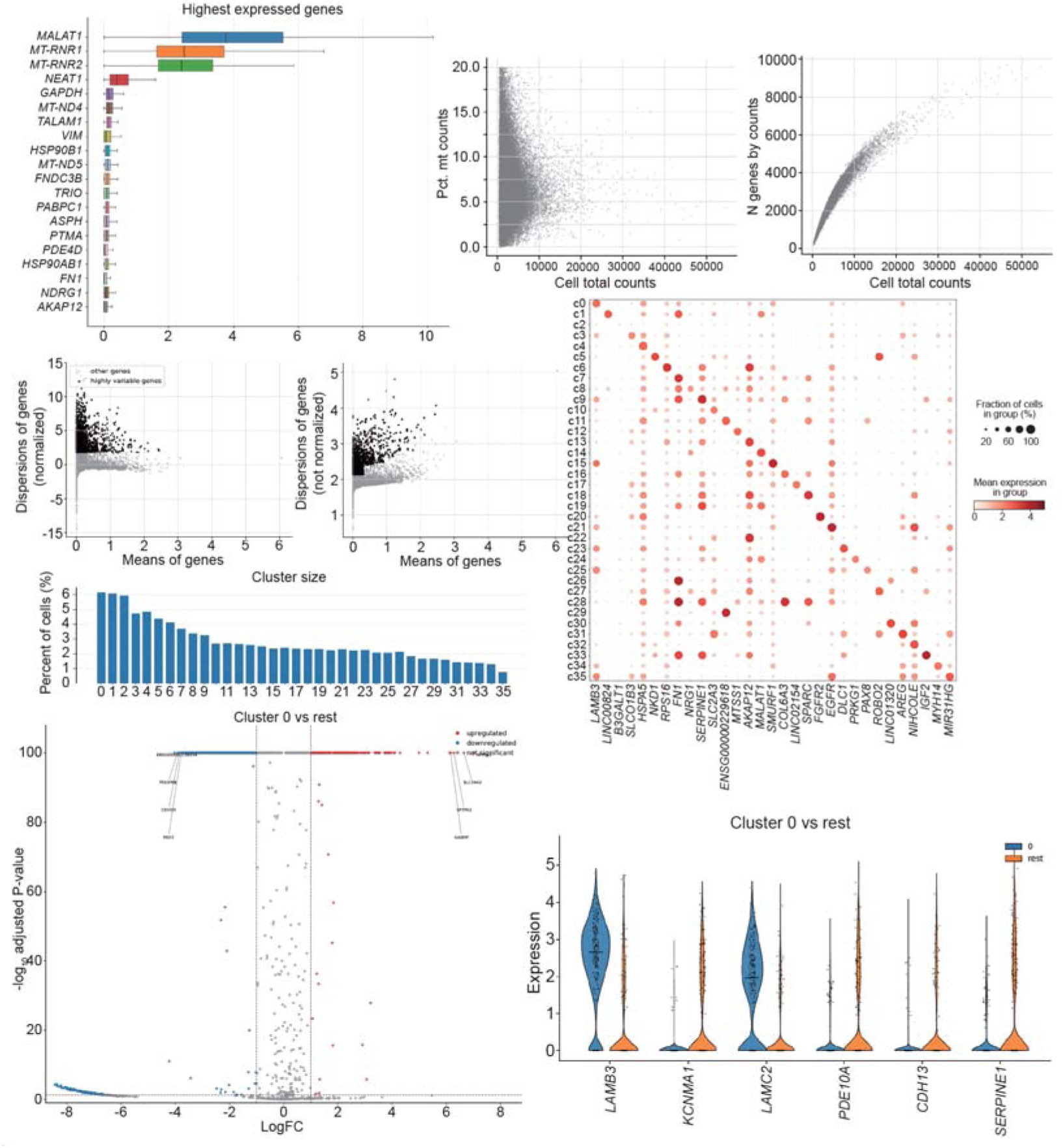
Representative exploratory outputs from the full 100M-cell Tahoe atlas. **(A)** highly expressed genes, **(B,C)** QC relationships between total counts, mitochondrial percentage and detected genes, **(D)** HVG selection before and after normalization, **(E)** marker dot plot, **(F)** Cluster composition, **(G)** differential-expression volcano plot, and **(H)** marker violin plots.

**Extended Data Figure 6.**
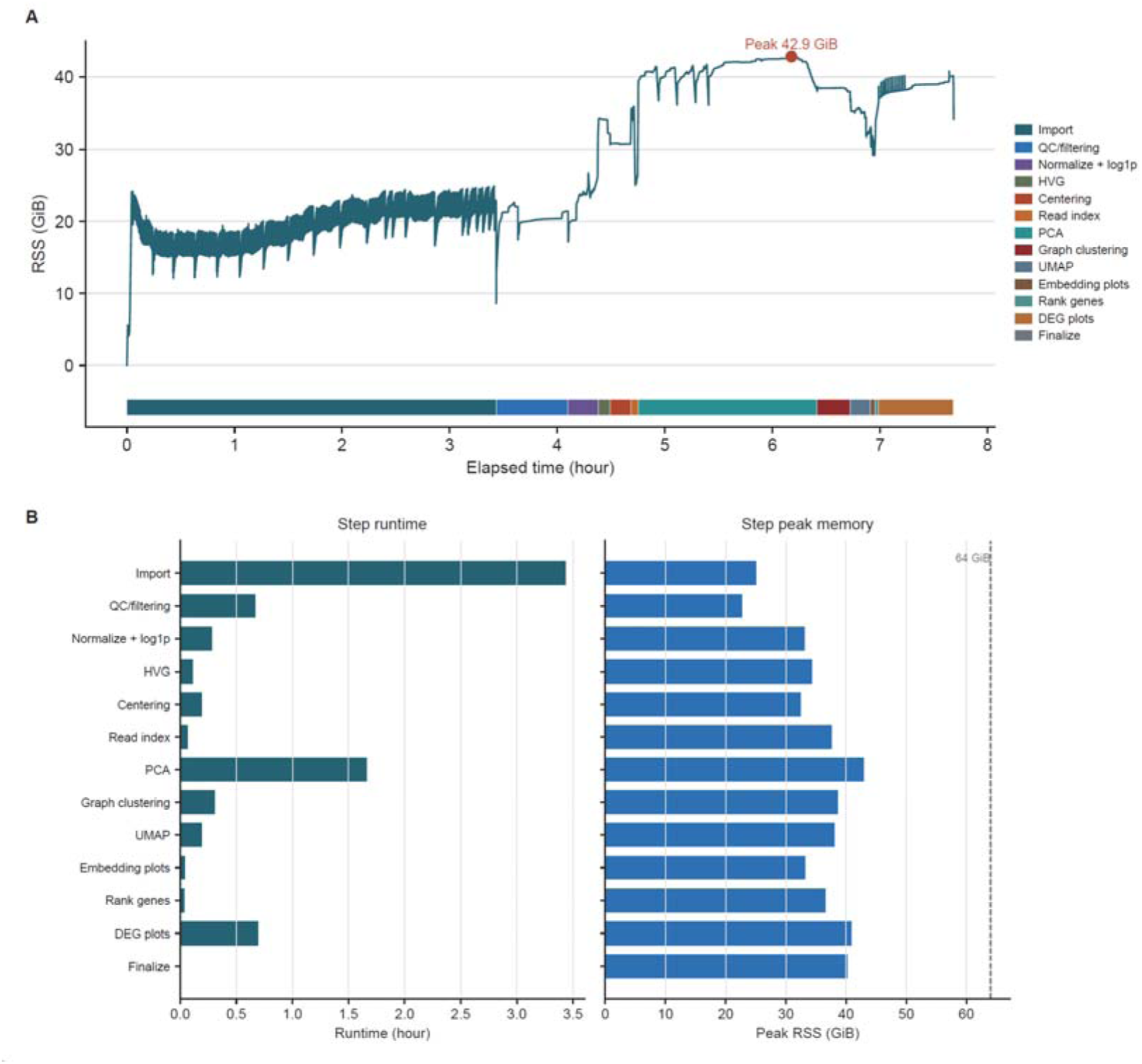
Resource profile of the 100M-cell Tahoe atlas analysis. A, RSS trace across the complete scAtlasPy workflow. Colored bars indicate individual workflow steps, and the peak RSS is annotated. **B,** Runtime (left) and peak RSS (right) for each workflow step. The dashed line indicates the 64 GiB system memory.

**Extended Data Figure 7.**
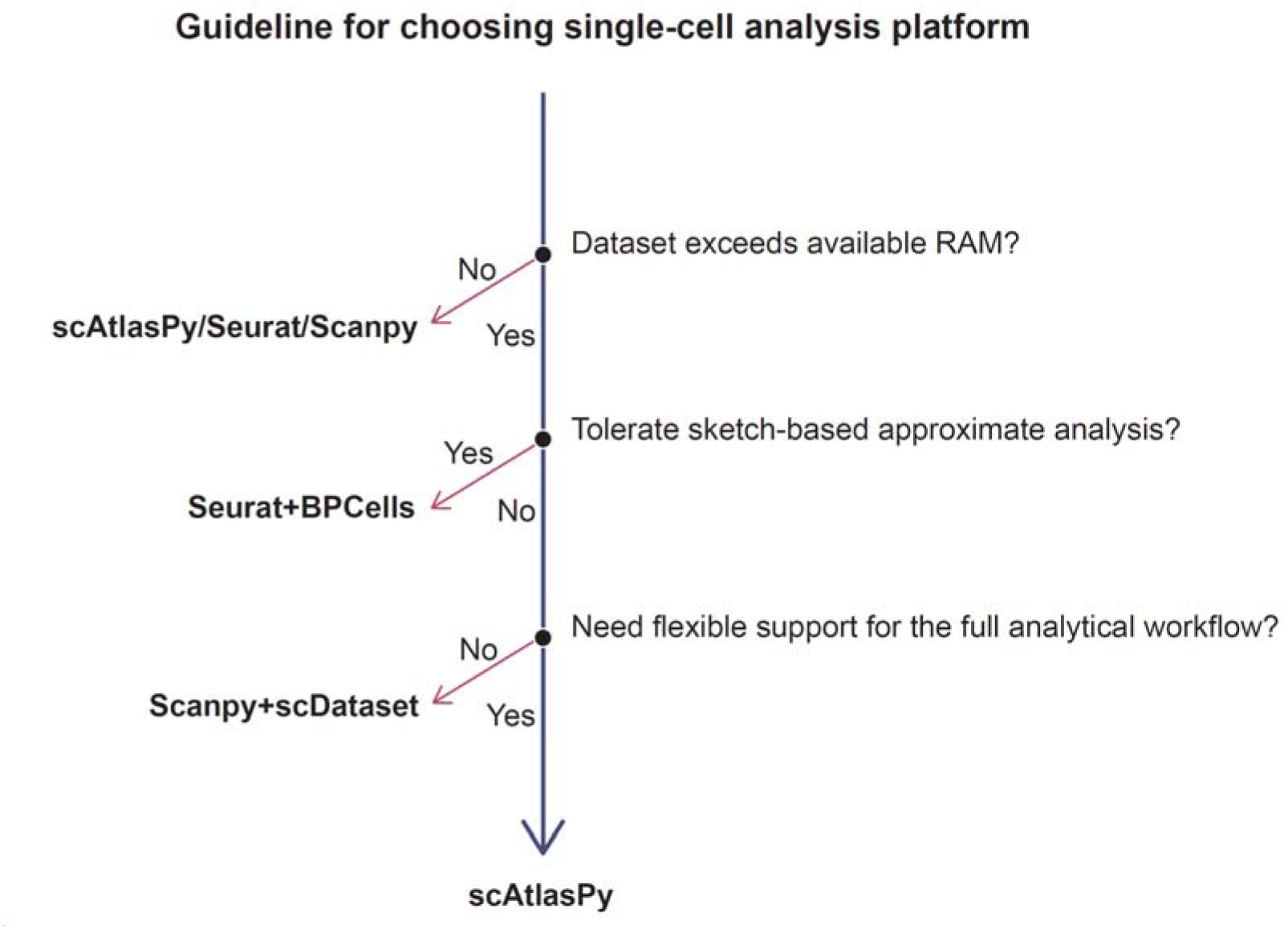
Guideline for choosing single-cell analysis platform.

## Extended Data Tables

**Extended Data Table 1.**
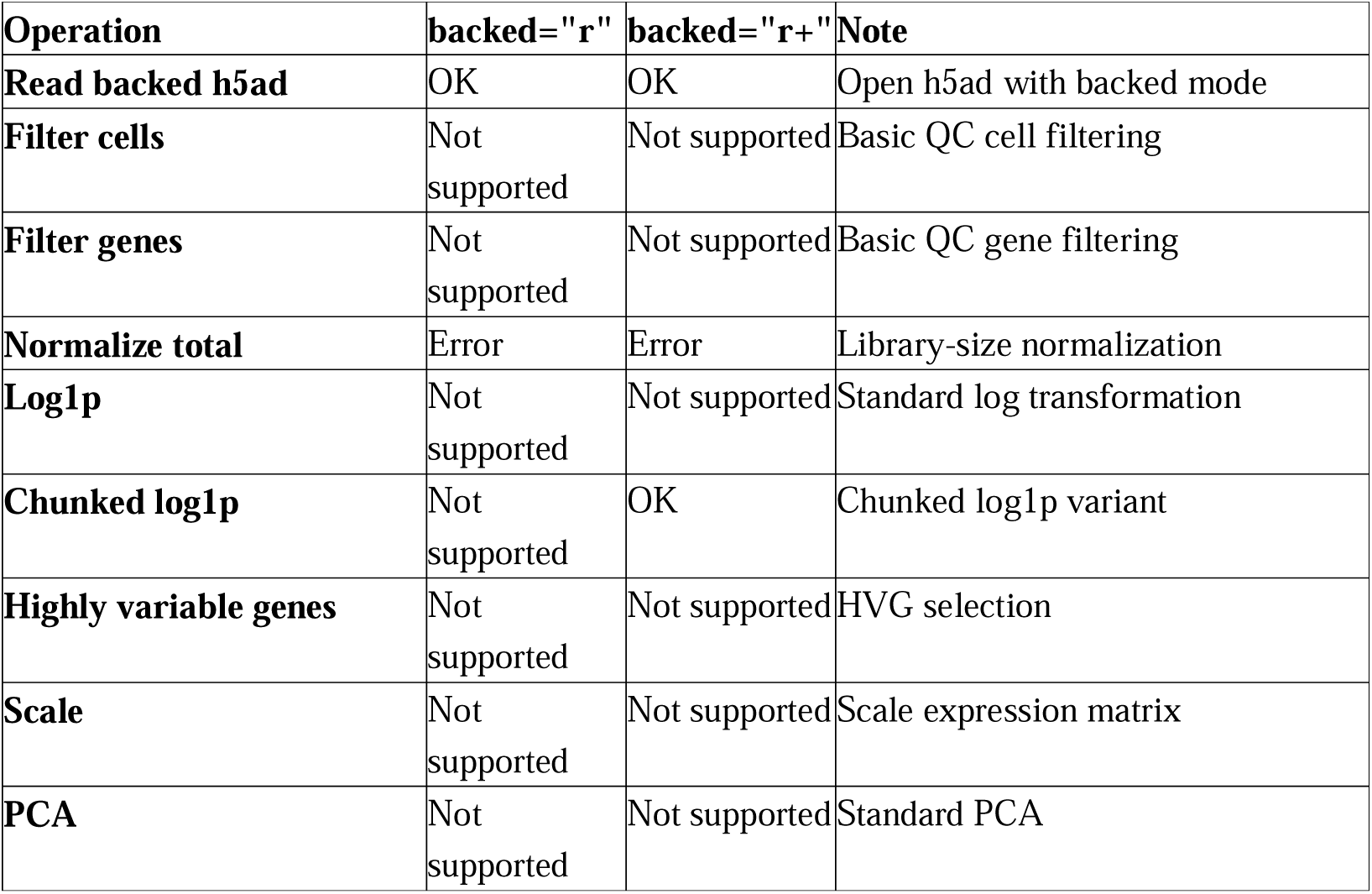
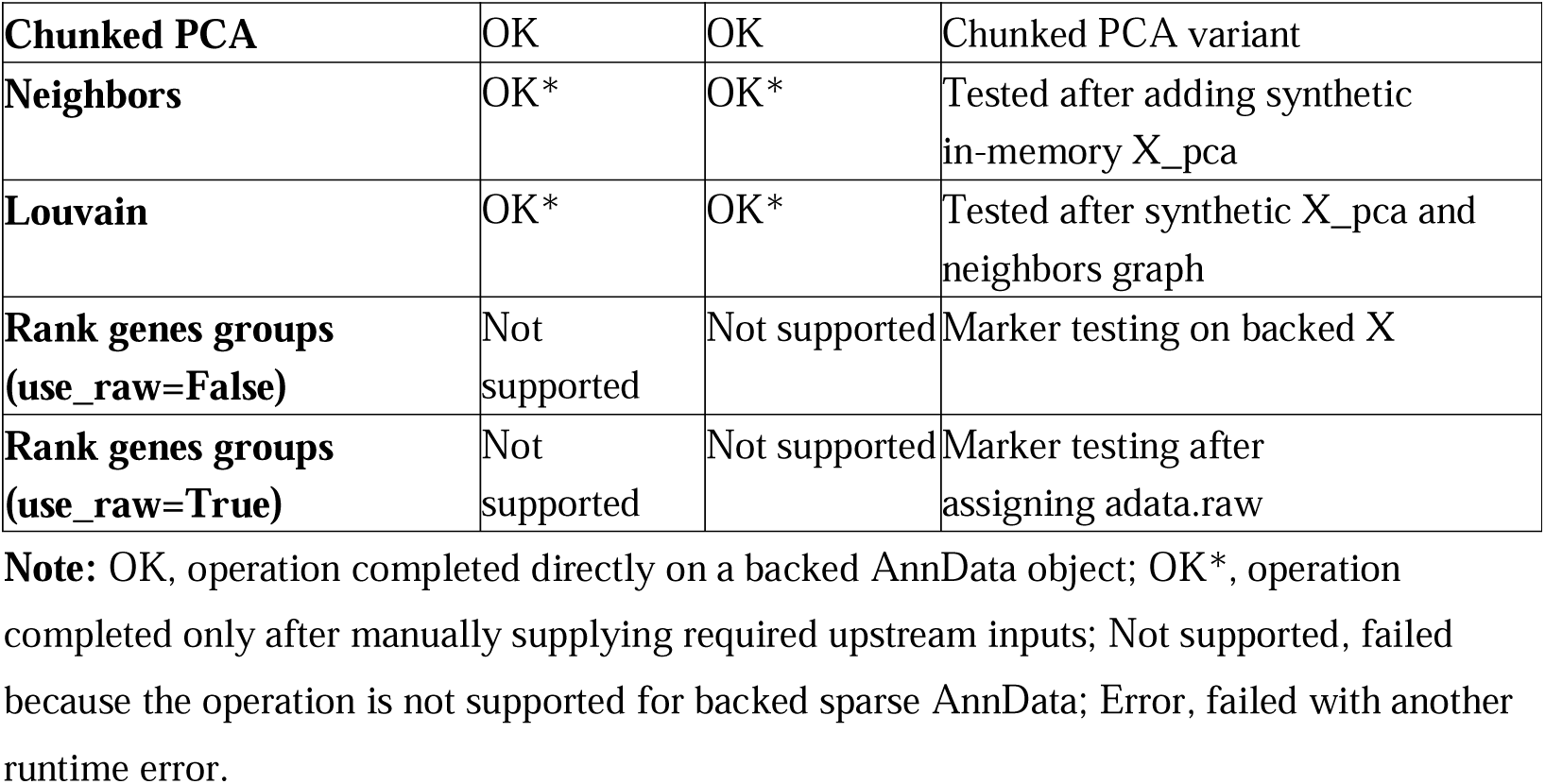
Support matrix of Scanpy backed AnnData for standard operations.

**Extended Data Table 2.**
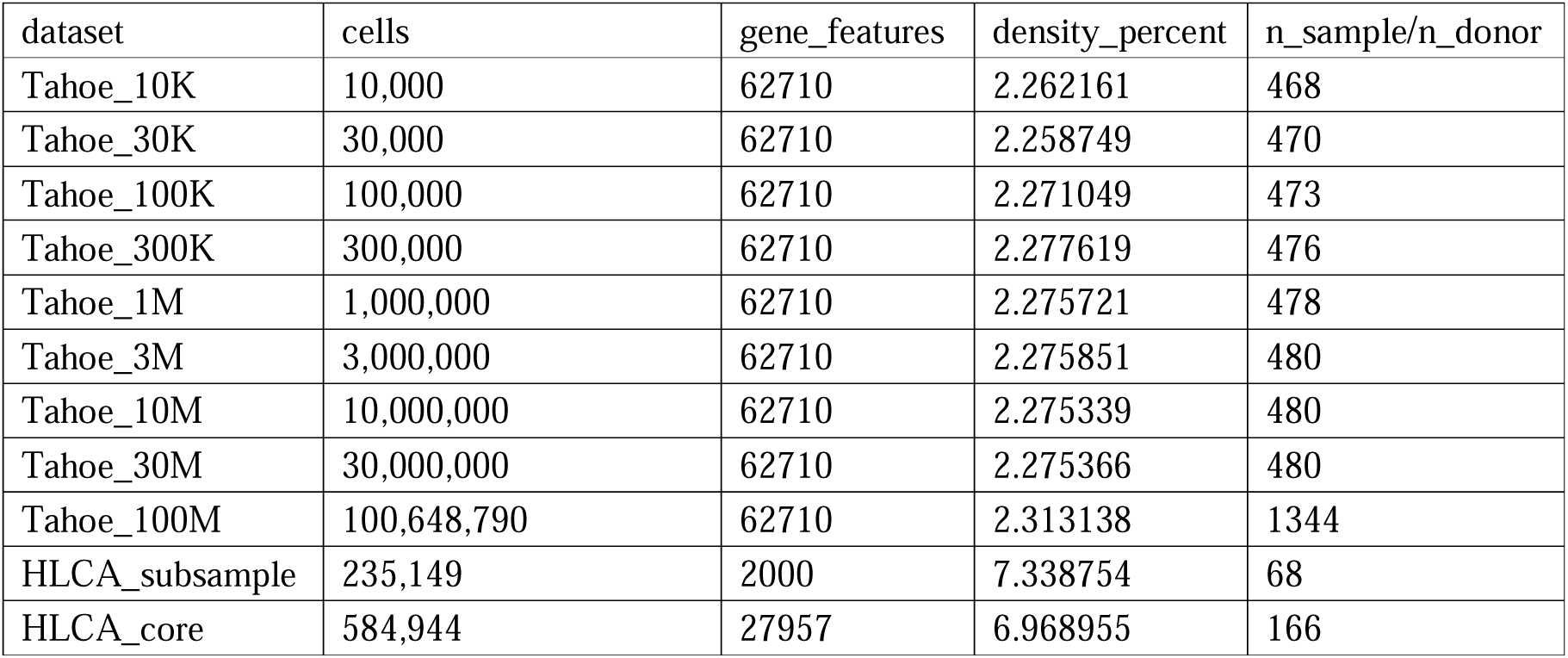
Summary of datasets used in this study.

**Extended Data Table 3.**
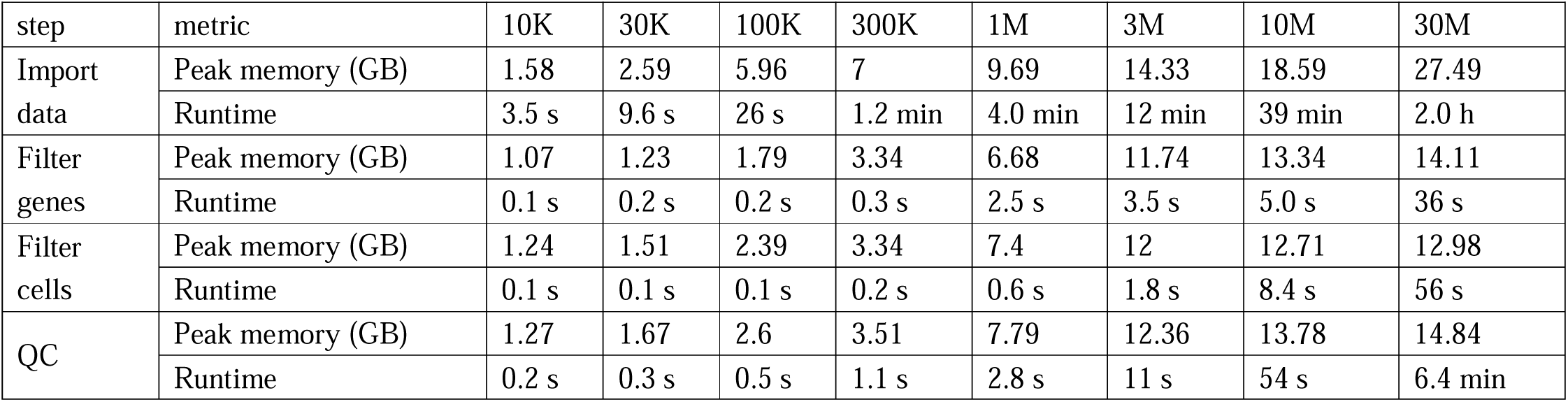
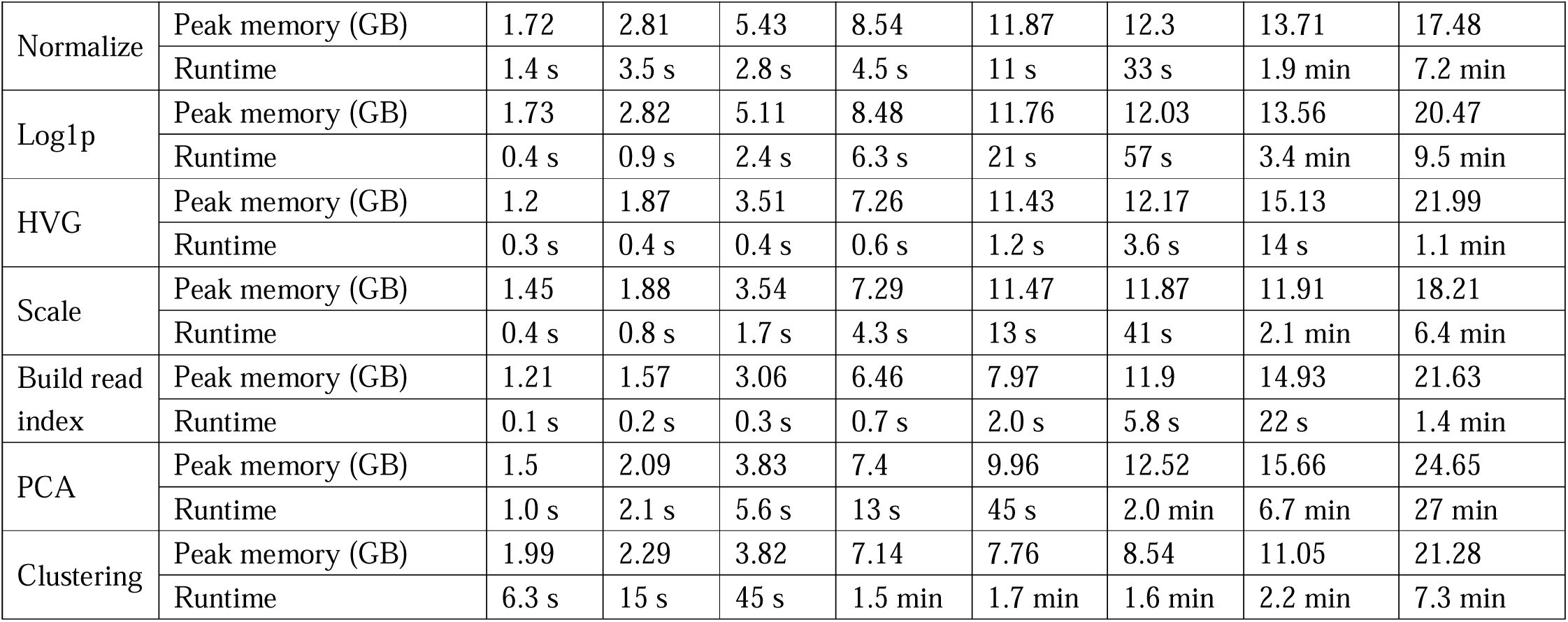
Step-level peak memory and runtime of scAtlasPy on scaling benchmarking tasks.

## References

1. Rood, J. E., Maartens, A., Hupalowska, A., Teichmann, S. A. & Regev, A. Impact of the Human Cell Atlas on medicine. Nature Medicine 28, 2486–2496 (2022).

2. Rood, J. E. et al. The Human Cell Atlas from a cell census to a unified foundation model. Nature 637, 1065–1071 (2025).

3. Hrovatin, K. et al. Considerations for building and using integrated single-cell atlases. Nature Methods 22, 41–57 (2025).

4. Zhang, J. et al. Tahoe-100M: a giga-scale single-cell perturbation atlas for context-dependent gene function and cellular modeling. bioRxiv 10.1101/2025.02.20.639398 (2025)

5. Wolf, F. A., Angerer, P. & Theis, F. J. SCANPY: large-scale single-cell gene expression data analysis. Genome Biology 19, 15 (2018).

6. Stuart, T. et al. Comprehensive integration of single-cell data. Cell 177, 1888–1902.e21 (2019).

7. Hao, Y. et al. Dictionary learning for integrative, multimodal and scalable single-cell analysis. Nature Biotechnology 42, 293–304 (2024).

8. Parks, B. & Greenleaf, W. Scalable high-performance single cell data analysis with BPCells. bioRxiv 10.1101/2025.03.27.645853 (2025)

9. D’Ascenzo, D. & Montesano, S. C. di. scDataset: scalable data loading for deep learning on large-scale single-cell omics. arxiv 10.48550/arXiv.2506.01883 (2026).

10. Pavan, K. & Saunders, A. AnnSQL: a Python SQL-based package for fast large-scale single-cell genomics analysis using minimal computational resources. Bioinformatics Advances 5, vbaf105 (2025).

11. Sikkema, L. et al. An integrated cell atlas of the lung in health and disease. Nature Medicine 29, 1563–1577 (2023).

12. Patikas, N. et al. Integration of large, complex single-cell datasets with Harmony2. bioRxiv 10.64898/2026.03.16.711825 (2026) doi:10.64898/2026.03.16.711825.

13. Virshup, I. et al. The scverse project provides a computational ecosystem for single-cell omics data analysis. Nature Biotechnology 41, 604–606 (2023).

14. Dicks, S., et al. GPU-accelerated single-cell analysis at scale with rapids-singlecell. arxiv 10.48550/arXiv.2603.02402 (2026)

15. Martinsson, P.-G. & Tropp, J. A. Randomized numerical linear algebra: Foundations and algorithms. Acta Numerica 29, 403–572 (2020).

16. Martinsson, P.-G. & Tropp, J. A. Randomized numerical linear algebra: Foundations and algorithms. Acta Numerica 29, 403–572 (2020).

