## extended data table 1~3, supplemental tables, supplemental information for "Atlas-scale single-cell analysis beyond in-memory paradigm with scAtlasPy": ex_table_1_scanpy_backed.docx

| **Operation** | **backed="r"** | **backed="r+"** | **Note** |
| --- | --- | --- | --- |
| **Read backed h5ad** | OK | OK | Open h5ad with backed mode |
| **Filter cells** | Not supported | Not supported | Basic QC cell filtering |
| **Filter genes** | Not supported | Not supported | Basic QC gene filtering |
| **Normalize total** | Error | Error | Library-size normalization |
| **Log1p** | Not supported | Not supported | Standard log transformation |
| **Chunked log1p** | Not supported | OK | Chunked log1p variant |
| **Highly variable genes** | Not supported | Not supported | HVG selection |
| **Scale** | Not supported | Not supported | Scale expression matrix |
| **PCA** | Not supported | Not supported | Standard PCA |
| **Chunked PCA** | OK | OK | Chunked PCA variant |
| **Neighbors** | OK* | OK* | Tested after adding synthetic in-memory X_pca |
| **Louvain** | OK* | OK* | Tested after synthetic X_pca and neighbors graph |
| **Rank genes groups (use_raw=False)** | Not supported | Not supported | Marker testing on backed X |
| **Rank genes groups (use_raw=True)** | Not supported | Not supported | Marker testing after assigning adata.raw |

Note:

OK, operation completed directly on a backed AnnData object; OK*, operation completed only after manually supplying required upstream inputs; Not supported, failed because the operation is not supported for backed sparse AnnData; Error, failed with another runtime error.
