## extended data table 1~3, supplemental tables, supplemental information for "Atlas-scale single-cell analysis beyond in-memory paradigm with scAtlasPy": ex_table_2_summary_of_datasets.docx

**Extended Data Table 2.** Summary of datasets used in this study

| dataset | cells | gene_features | density_percent | n_sample/n_donor |
| --- | --- | --- | --- | --- |
| Tahoe_10K | 10,000 | 62710 | 2.262161 | 468 |
| Tahoe_30K | 30,000 | 62710 | 2.258749 | 470 |
| Tahoe_100K | 100,000 | 62710 | 2.271049 | 473 |
| Tahoe_300K | 300,000 | 62710 | 2.277619 | 476 |
| Tahoe_1M | 1,000,000 | 62710 | 2.275721 | 478 |
| Tahoe_3M | 3,000,000 | 62710 | 2.275851 | 480 |
| Tahoe_10M | 10,000,000 | 62710 | 2.275339 | 480 |
| Tahoe_30M | 30,000,000 | 62710 | 2.275366 | 480 |
| Tahoe_100M | 100,648,790 | 62710 | 2.313138 | 1344 |
| HLCA_subsample | 235,149 | 2000 | 7.338754 | 68 |
| HLCA_core | 584,944 | 27957 | 6.968955 | 166 |
