## extended data table 1~3, supplemental tables, supplemental information for "Atlas-scale single-cell analysis beyond in-memory paradigm with scAtlasPy": ex_table_3_step_level_mem_and_time.docx

| step | metric | 10K | 30K | 100K | 300K | 1M | 3M | 10M | 30M |
| --- | --- | --- | --- | --- | --- | --- | --- | --- | --- |
| Import data | Peak memory (GB) | 1.58 | 2.59 | 5.96 | 7 | 9.69 | 14.33 | 18.59 | 27.49 |
|  | Runtime | 3.5 s | 9.6 s | 26 s | 1.2 min | 4.0 min | 12 min | 39 min | 2.0 h |
| Filter genes | Peak memory (GB) | 1.07 | 1.23 | 1.79 | 3.34 | 6.68 | 11.74 | 13.34 | 14.11 |
|  | Runtime | 0.1 s | 0.2 s | 0.2 s | 0.3 s | 2.5 s | 3.5 s | 5.0 s | 36 s |
| Filter cells | Peak memory (GB) | 1.24 | 1.51 | 2.39 | 3.34 | 7.4 | 12 | 12.71 | 12.98 |
|  | Runtime | 0.1 s | 0.1 s | 0.1 s | 0.2 s | 0.6 s | 1.8 s | 8.4 s | 56 s |
| QC | Peak memory (GB) | 1.27 | 1.67 | 2.6 | 3.51 | 7.79 | 12.36 | 13.78 | 14.84 |
|  | Runtime | 0.2 s | 0.3 s | 0.5 s | 1.1 s | 2.8 s | 11 s | 54 s | 6.4 min |
| Normalize | Peak memory (GB) | 1.72 | 2.81 | 5.43 | 8.54 | 11.87 | 12.3 | 13.71 | 17.48 |
|  | Runtime | 1.4 s | 3.5 s | 2.8 s | 4.5 s | 11 s | 33 s | 1.9 min | 7.2 min |
| Log1p | Peak memory (GB) | 1.73 | 2.82 | 5.11 | 8.48 | 11.76 | 12.03 | 13.56 | 20.47 |
|  | Runtime | 0.4 s | 0.9 s | 2.4 s | 6.3 s | 21 s | 57 s | 3.4 min | 9.5 min |
| HVG | Peak memory (GB) | 1.2 | 1.87 | 3.51 | 7.26 | 11.43 | 12.17 | 15.13 | 21.99 |
|  | Runtime | 0.3 s | 0.4 s | 0.4 s | 0.6 s | 1.2 s | 3.6 s | 14 s | 1.1 min |
| Scale | Peak memory (GB) | 1.45 | 1.88 | 3.54 | 7.29 | 11.47 | 11.87 | 11.91 | 18.21 |
|  | Runtime | 0.4 s | 0.8 s | 1.7 s | 4.3 s | 13 s | 41 s | 2.1 min | 6.4 min |
| Build read index | Peak memory (GB) | 1.21 | 1.57 | 3.06 | 6.46 | 7.97 | 11.9 | 14.93 | 21.63 |
|  | Runtime | 0.1 s | 0.2 s | 0.3 s | 0.7 s | 2.0 s | 5.8 s | 22 s | 1.4 min |
| PCA | Peak memory (GB) | 1.5 | 2.09 | 3.83 | 7.4 | 9.96 | 12.52 | 15.66 | 24.65 |
|  | Runtime | 1.0 s | 2.1 s | 5.6 s | 13 s | 45 s | 2.0 min | 6.7 min | 27 min |
| Clustering | Peak memory (GB) | 1.99 | 2.29 | 3.82 | 7.14 | 7.76 | 8.54 | 11.05 | 21.28 |
|  | Runtime | 6.3 s | 15 s | 45 s | 1.5 min | 1.7 min | 1.6 min | 2.2 min | 7.3 min |
