## extended data table 1~3, supplemental tables, supplemental information for "Atlas-scale single-cell analysis beyond in-memory paradigm with scAtlasPy": Supplementary Information.docx

Supplementary Note 1

**Large input matrix.**

We observed the architectural limitation when Seurat was applied directly to a large input expression matrix in the scaling benchmark. In this experiment, we used the Tahoe down-sampled 3M-cell dataset, stored as tahoe100M_plate1_5_subsample_3000000_cells.h5ad. The benchmark attempted to import the full sparse expression matrix into the traditional Seurat workflow before any downstream filtering, normalization, dimensionality reduction, or clustering steps were performed.

Seurat failed during the import/access step. The run terminated after 264.16 seconds with a peak RSS memory of 54.34 GB. The recorded error was:

simpleError: SVT_SparseMatrix object contains too many nonzero values (4281558519) to "fit" in a CsparseMatrix derivative

The reported number of nonzero entries was 4,281,558,519, which exceeds the 2^31 - 1 limit imposed by the CsparseMatrix representation used by the R Matrix infrastructure. Because the failure occurred during the initial construction/conversion of the input sparse matrix, all downstream steps, including cell filtering, gene filtering, QC, normalization, HVG selection, scaling, PCA, and clustering, were skipped.

**Large intermediate matrix.**

We evaluated Seurat integration on a preprocessed subset of the HLCA core dataset, stored as HLCA_core__subsample__hvg_2000.h5ad. The dataset contained 235,149 cells and 2,000 highly variable genes after down-sampling, log transformation, and HVG selection. The expression matrix was stored as a sparse CSR matrix with 34,514,013 nonzero entries.

We imported the dataset into Seurat 5.5.0 with Matrix 1.7.5, used donor_id as the batch key, and split the data into 44 donor-level batches. Since the input matrix was already log-normalized and restricted to HVGs, we skipped additional normalization and used all 2,000 genes as integration features. We then ran the standard Seurat integration workflow: constructing Seurat objects, splitting by donor, finding pairwise integration anchors with FindIntegrationAnchors(), and integrating the batches with IntegrateData().

Seurat successfully completed pairwise anchor detection for all 946 donor-pair comparisons. The failure occurred later during IntegrateData(), while merging larger groups of donor batches and computing the integration matrix. The run terminated inside Seurat’s FindIntegrationMatrix step with the following Matrix error:

“

Error in ..subscript.2ary(x, l[[1L]], l[[2L]], drop = drop[1L]) :

x[i,j] too dense for [CR]sparseMatrix; would have more than 2^31-1 nonzero entries

Calls: IntegrateData ... FindIntegrationMatrix -> [ -> [ -> .subscript.2ary -> ..subscript.2ary

”

This case demonstrates that the failure was not only caused by the size of the input expression matrix itself but also can raise by the intermediate matrix in the Seurat workflow.This run used approximately 113 GB peak memory and failed after about 3.5 hours. The log files of these runs and the scripts to reproduce this issue are available in our repository.

Similar issues were also previously discussed in the GitHub of Seurat.

<https://github.com/satijalab/seurat/issues/3650>

<https://github.com/satijalab/seurat/issues/4621>

Supplementary Note 2

Scanpy performs on-disk analysis by linking H5AD files to backed AnnData objects. We used a small dataset to test whether backed AnnData supports individual basic operations. We conducted an operation-level support test to identify which steps of a standard Scanpy workflow can be executed directly on backed AnnData objects. This experiment was designed as a supplementary diagnostic rather than a performance benchmark. We used the smallest Tahoe100M plate1–5 subsample, which contains 10,000 cells, 62,710 genes, and 14,186,011 non-zero expression values. The input file was loaded using sc.read_h5ad(..., backed=...), and each operation was tested under both backed="r" and backed="r+" modes.

Each operation was evaluated independently in an isolated child process using probe_scanpy_backed_step_support.py. No timeout threshold was applied, as delayed or slow completion was regarded as inherent to the practical behavior of backed-mode execution. For operations that normally depend on upstream results, we provided only the minimal prerequisites required to isolate support for the target operation. For instance, neighbor graph construction was tested after injecting a synthetic in-memory X_pca embedding, and Leiden clustering was tested after constructing a synthetic neighbor graph. These results therefore indicate whether the tested function can operate on a backed AnnData object when its dependencies are satisfied, rather than whether the full end-to-end backed-mode pipeline can reach that step.

The resulting support matrix is summarized in Extended Data Table 1. In this table, OK indicates that the operation completed successfully when applied directly to a backed AnnData object. OK* indicates successful execution only after manually supplying required upstream inputs, such as a PCA embedding or neighbor graph. Not supported indicates failure due to an explicit Scanpy or AnnData limitation for backed sparse data. Error indicates an unclassified runtime failure that did not produce an explicit backed-mode unsupported-operation message.

Our results demonstrate that backed AnnData objects can be opened successfully in both read-only (r) and read–write (r+) modes. However, many core preprocessing operations are not supported on backed sparse matrices. Cell filtering, gene filtering, standard log1p transformation, scaling, standard PCA, and differential expression testing all failed with explicit unsupported-operation errors for backed sparse matrices. Highly variable gene selection also failed because the backed sparse format does not support the required copy operation. Chunked log1p succeeded only in r+ mode, whereas chunked PCA succeeded in both r and r+ modes. Neighbor graph construction and Leiden clustering succeeded when synthetic upstream representations were provided, but these results do not imply that a complete end-to-end backed-mode preprocessing pipeline is executable. Collectively, the support matrix reveals that Scanpy backed mode enables convenient file-backed data access with limited operation support, but does not enable the full standard preprocessing, clustering, and differential expression workflow directly on backed sparse H5AD data.
